# States and traits of neural irregularity in the age-varying human brain

**DOI:** 10.1101/103432

**Authors:** Leonhard Waschke, Malte Wöestmann, Jonas Obleser

## Abstract

Sensory representations of the physical world and thus human percepts are susceptible to fluctuations in brain state or “neural irregularity”. Furthermore, aging brains display altered levels of irregularity. We here show that a single, within-trial information-theoretic measure (weighted permutation entropy) captures neural irregularity in the human electroencephalogram as a proxy for both, trait-like differences between individuals of varying age, and state-like fluctuations that bias perceptual decisions. First, the overall level of neural irregularity increased with participants‘ age, paralleled by a decrease in variability over time, likely indexing age-related disintegration on structural and functional levels of brain activity. Second, states of higher neural irregularity were associated with optimized sensory encoding and a subsequently increased probability of choosing the first of two physically identical stimuli. In sum, neural irregularity not only characterizes behaviorally relevant brain states, but also can identify trait-like changes that come with age.

## Introduction

Brain activity, no matter if ongoing or evoked by sensory stimulation, is inherently variable or irregular—a phenomenon often termed “neural noise” (Faisal, Selen, and Wolpert 2008; Arieli et al. 1996). The degree of this neural irregularity in neocortex is thought to track perceptual sensitivity (Arazi, Censor, and Dinstein 2017), but might also capture more lasting changes in brain activity that come with age. Moreover, neural irregularity is discussed as a potential marker of different neuropsychiatric disorders (Dinstein, Heeger, and Behrmann 2015). What has been missing so far, however, is a feasible and versatile metric of neural irregularity in the human EEG that would capture both, the trait-like, across-participants variations in neural irregularity and state-like, across-trials variations within the individual that co-determine the perceptual decisions we take facing ambiguous or missing physical evidence.

Across individuals, older age is accompanied by decreasing trial-to-trial brain signal variability (Grady and Garrett 2014; Garrett et al. 2013) and increasing 1/f noise in the electroencephalogram (EEG) (i.e., flatter EEG frequency spectra; Sleimen-Malkoun et al., 2015; Voytek et al., 2015). Those changes likely reflect a shift from distributed to more local processing. Spontaneous brain activity, and in particular its variability, thus tracks stable, trait-like differences *between* individuals of varying age.

*Within* an individual, the (ir-)regularity of ongoing cortical activity might be a constituent of brain states that influence sensory encoding, perception, and ensuing behavior. For instance, the neural response to a sensory stimulus depends on a participant’s cortical activity preceding stimulus onset (Arieli et al. 1996). More precisely, encoding and representing sensory information is more thorough in the presence of a desynchronized (irregular) state as compared to a synchronized (regular) state (Marguet and Harris 2011; Pachitariu et al. 2015). Intra-individual variability of spontaneous cortical activity hence represents fluctuations of cortical states that entail perceptual and behavioral relevance.

But how can we track neural irregularity simultaneously on the group-level and on the individual level? A desirable measure of neural irregularity should not only predict differences in encoding and perceptual decisions between identical stimuli (in terms of a neural state), but also track changes in the degree of irregularity that come with older age (in terms of a neural trait) (Arieli et al. 1996). The most promising measures in this respect, offering sufficient resolution in time and space, derive from information theory: Entropy measures have been used extensively to research epileptic seizures (Nicolaou and Georgiou 2012; Dickten et al. 2016) or vigilance states (Bruzzo et al. 2008) and cognitive processes in the EEG (O’Hara et al., 2013). Information-theoretic measures of single trial time-courses should allow tracking whether neural irregularity (i) can be linked to perceptual decisions, reflecting trial-by-trial brain-state changes and (ii) can serve as a trait-like marker for individual age-related change in brain activity. Here, we demonstrate that these requirements are met by a single-trial, time-resolved, ordinal measure of entropy, called Weighted Permutation Entropy (WPE, Fadlallah et al., 2013).

As a test case, the present study aimed at predicting participants’ perceived pitch differences between two identical tones from neural irregularity. Computational modeling (Micheyl, McDermott, and Oxenham 2009) as well as work on the auditory evoked potential (Bernasconi et al. 2011) brought to bear the hypothesis that fluctuations in spontaneous neural activity are the most likely source for varying percepts in response to identical stimuli. Thus, if an individual reports perceptual differences for identical stimuli, these reports are likely influenced by the momentary or pre-stimulus degree of irregularity in the perceiver’s neural system.

We demonstrate that the irregularity of the scalp EEG response, operationalized as pre­ stimulus WPE, offers a clear relationship to spontaneous, pre-stimulus brain states as often described in terms of neural oscillatory power;to the sensory fidelity with which a stimulus is neurally encoded; and to the ensuing perceptual decision upon this stimulus. Moreover, entropy proves to be a sensitive marker for inter-individual, trait-like changes of neural irregularity that come with age.

## Results

Human participants of varying age (*n* = 19; 19–74 years; 49 ± 18.75 Median ± Semi-lnterquartile range, sIQR) were asked to compare the pitches of two identical pure tones. We were interested in (i) the influence of pre-stimulus irregularity in ongoing EEG activity on sensory encoding and the impending perceptual decision within individuals, as well as (ii) how these patterns of neural irregularity differ across individuals and change with age.

While participants were able to perform perceptual decisions on these physically identical tones and indeed did perceive pitch differences (see supplementary figures S1 and S2; Amitay et al., 2006; Micheyl et al., 2009), decisions for the first tone (S1, 57% on average) were overall slightly more frequent than for the second one (S2, 43% on average; 13 ± 4.5% ± SEM for the difference; *t*_18_ = 3.0, *p* = .01, *r_e_* =.58), representing a slight bias towards perceiving S1 as higher in pitch.

## Patterns of irregularity in ongoing activity change with age

We investigated both interindividual differences, or traits, as well as intraindividual variation, or states, in pre-stimulus entropy: First, we compared the variability of entropy in a pre-stimulus time­ window (–.4 to –.1 s) across trials and participants using an lntraclass-correlation coefficient (ICC, ranging from 0 to 1, Fig. 1B, upper panel). The ICC allows us quantify to which degree total variance is due to differences between participants.

**Figure 1.**
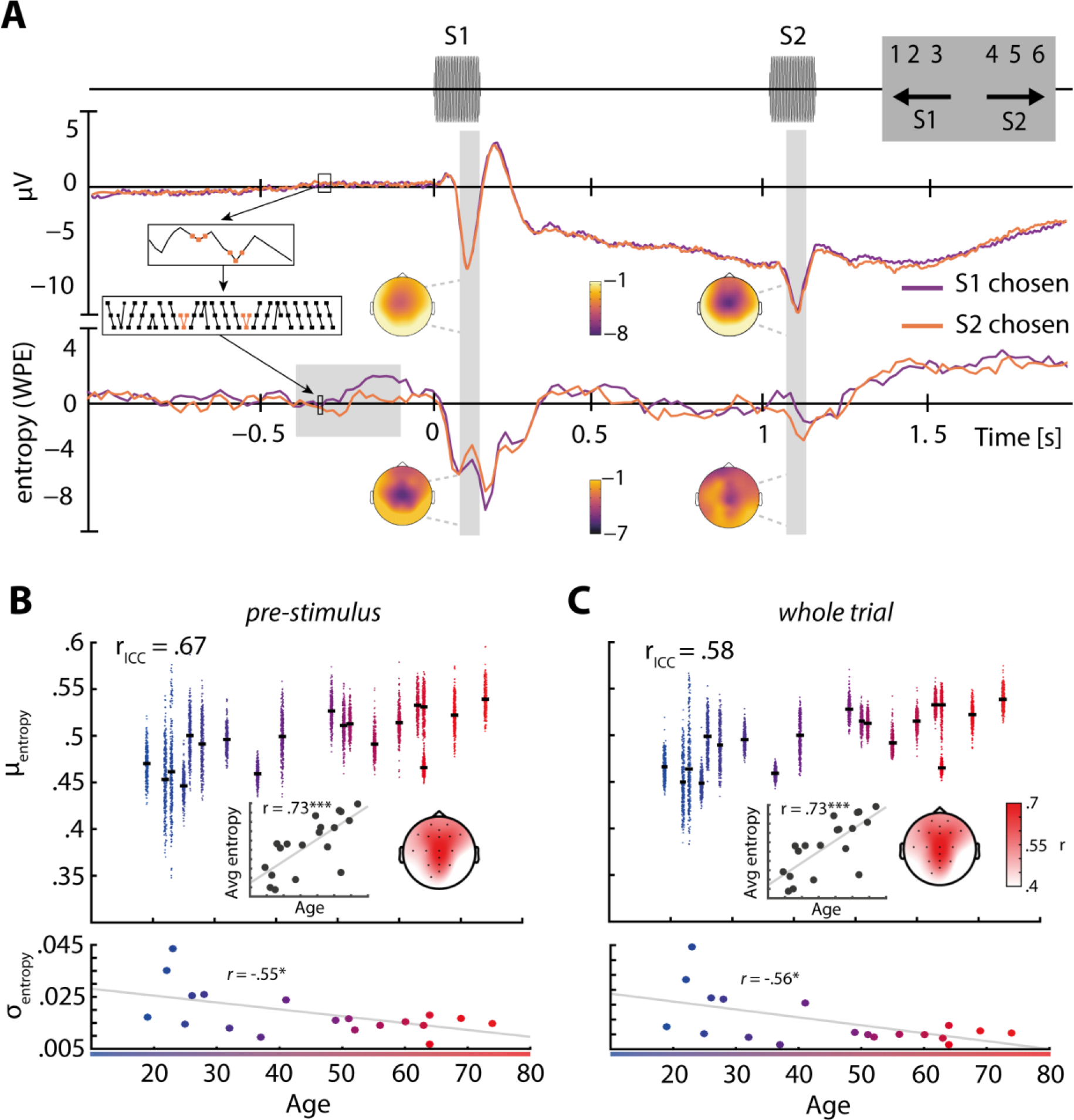
Trial structure, evoked responses, and entropy across age. A, Trial structure and average responses. Two 650Hz pure tones were presented separated by a fixed lSI of 900 ms and followed by the response screen (top panel). The event-related potential (ERP) at electrode Cz (baseline –.5 to 0 s) is shown for trials in favor of the first (S1, purple) and the second tone (S2, orange) in the middle panel. Topographies correspond to the N1 time-windows after S1 and S2 (grey areas). Colors show average microvolt values. Entropy (WPE) is computed by taking small snippets of voltage data, transforming them to rank sequences and computing the entropy of their frequency of occurrence. This procedure is illustrated in the small inset boxes that connect ERP and WPE representations. In the bottom panel, average WPE time courses of both conditions are shown (baselined relative to −1.5 to −1 s). Colors on topographies correspond to average WPE. **B, Pre-stimulus entropy (WPE) across age.** Trial-wise averages of entropy in a pre-stimulus time window (–.4 to –.1 s, electrode Cz; grey shaded area) are shown as clouds of dots per subject (upper panel, warmer colors code higher age), black bars indicate average entropy across trials. Average entropy increased with age (upper panel, inset electrode Cz), colors on topography correspond to average correlation coefficients r. The standard deviation of entropy across trials decreased with age (lower panel). **C**, Same as B but for entropy data from whole trials (i.e., −2 to 2 s).*p < .05;*** p < .001

As indicated by a fairly high ICC (*r_ICC_* = .67; Cicchetti, 1994), the sizable within-subject, trial-to­ trial variability was in fact clustered at the level of participants (Fig. 1B). Thus, the variation in entropy values observed overall is attributable to both, state-like trial-by-trial fluctuation and trait-like between-subjects variation.

Exploring this between-subjects variation of entropy, we correlated participants’ age with their individual pre-stimulus entropy averaged across trials (Fig. 1B, upper panel, inset), and their corresponding intra-individual variation (standard deviation; Fig. 1B, lower panel). Participants’ age exhibited a high positive correlation with average pre-stimulus entropy (*r* = .73, *p* = .0004). This correlation, shown for electrode Cz in Fig. 1B, was significant at a large set of fronto-central and parietal electrodes (Fig. 1B, topography, highlighted electrodes *p* ≤ .05) but clustered towards central electrodes, forming a topography reminiscent of auditory evokes responses. Notably, the standard deviation of pre-stimulus entropy across trials was negatively correlated with age (*r* = −.55, *p* = .02, Fig. 1B, lower panel). Older participants thus exhibit increased levels of average pre-stimulus entropy, while at the same time showing decreased trial-to-trial variability of pre-stimulus entropy.

Note that when repeating both analyses based on mean entropy across the whole trial instead (−2 to 2 s), the observed patterns were remarkably stable (Fig. 1C, see also Fig. S3). This underlines the robustness of the weighted permutation entropy measure against overly specific analysis choices when serving as a trait-like marker of neural irregularity.

## Irregularity of the EEG signal approximates 1/f noise

To validate the increase of pre-stimulus entropy with age, we related it to an integrative measure of spectral shape, power spectral density (PSD): A fitted linear slope to the PSD is usually negative and reflects the inverse power-law dynamics of the human EEG (Voytek et al. 2015). Expectedly, individual PSD slopes became less negative with increasing participant age (*r* = .80, *p* = 4 × 10^−5^; Fig. 2A), with decreased power at low and increased power at higher frequencies, respectively. Notably, a similar effect could be substantiated when correlating age with the PSD slope derived from higher frequencies (30–60 Hz; *r* = .62, *p* = .001; Fig. S4) recently suggested to capture changes in excitation inhibition (E/1) balance (Gao, Peterson, and Voytek 2016).

**Figure 2.**
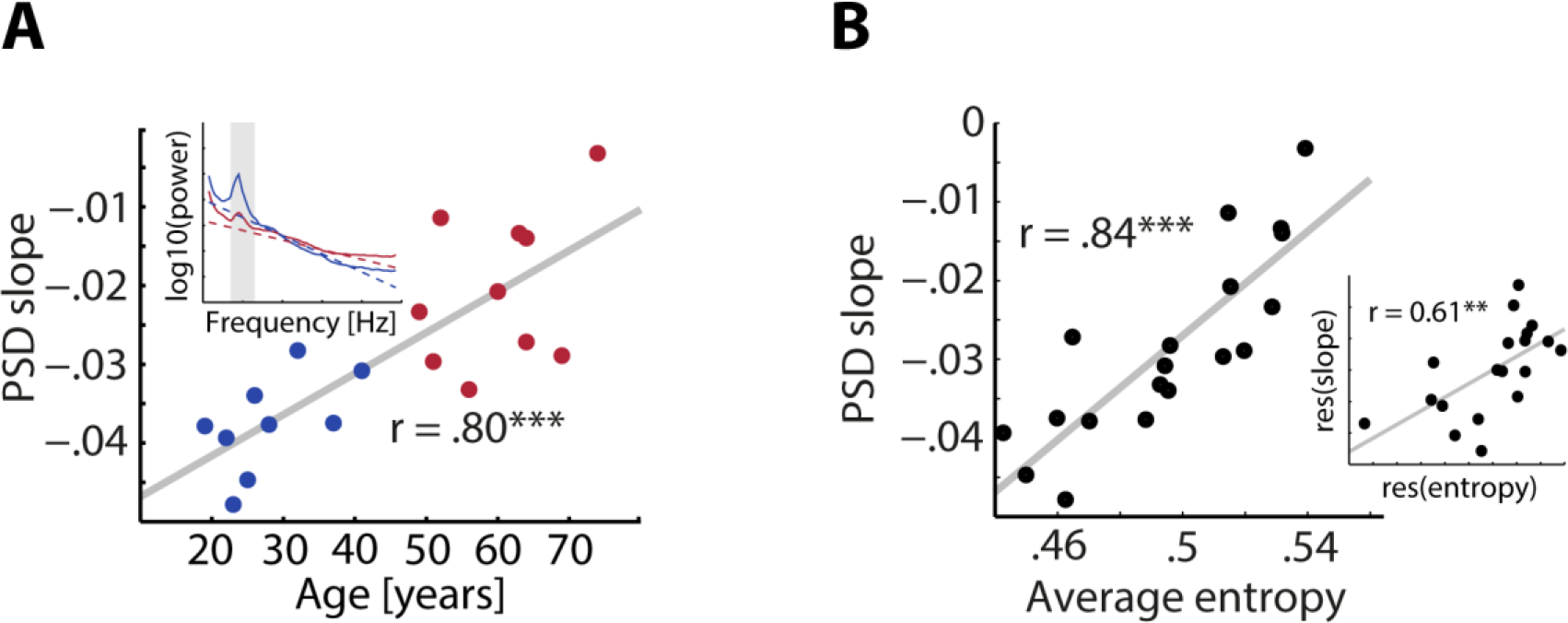
Weighted permutation entropy captures 1/f noise. A, PSD slope changes with age. The slope of the PSD (inset) at electrode Cz over frequencies 1–30 Hz becomes less negative with increasing age. Younger subjects are shown in blue and older subjects in red. **B, WPE and PSD slopes are linearly related.** The correlation of average entropy (WPE) and PSD slope is shown. The inset shows the same correlation after regressing out age.** *p* < .01;*** *p* < .0001.

Since both average entropy and PSD slopes potentially capture a common underlying mechanism, the overall degree of irregularity, we correlated both measures and found a strikingly linear relationship (*r* = .84, *p* = 8 × 10^−6^ Fig. 2B). Importantly, this relationship of the time-domain WPE measure and the spectral-domain PSD measure of neural irregularity retained a substantial effect size even after contraIIing for age (*r* = .61, *p* = .006, Fig. 2B, inset). This validates the utility of entropy and PSD slope a s trait-like markers of interindividual variability above and beyond senescent change in the time and frequency domain, respectively.

## Pre-stimulus irregularity captures low-frequency power and peri-stimulus phase alignment

Previous work in non-human animals has characterized differences in synchrony and power of low-frequency oscillations that influence encoding and subsequent representations of stimuli (Pachitariu et al. 2015; Marguet and Harris 2011). Thus, we aimed to validate pre-stimulus entropy as a measure of neural state by exploring its potency to also approximate fluctuations in low-frequency power and phase coherence of the human EEG.

To this end, we binned trials by normalized, pre-stimulus entropy (–.4 to –.1 s) into four bins of equal trial number before calculating average power and inter-trial phase coherence (ITPC) in each bin (cf.bar graphs in Fig. 3). Separately for each participant we then fitted linear slopes to average power and ITPC, respectively, across the four bins of increasing pre-stimulus entropy. A cluster-based permutation test comparing slopes against zero revealed a broad negative cluster of changing oscillatory power with increasing entropy (–.85 to .25 s, 1–29 Hz) displaying a fronto-central topography (Fig. 3A, *p* = .0009, *R* = –.71). Notably, the same analysis for EEG power in higher frequencies (30–70Hz) did not reveal any significant change across bins (all *p* > .05, Fig. S5). Thus, pre-stimulus entropy tracked changes predominantly in low-frequency power, with lower power accompanying higher time-domain irregularity.

**Figure 3.**
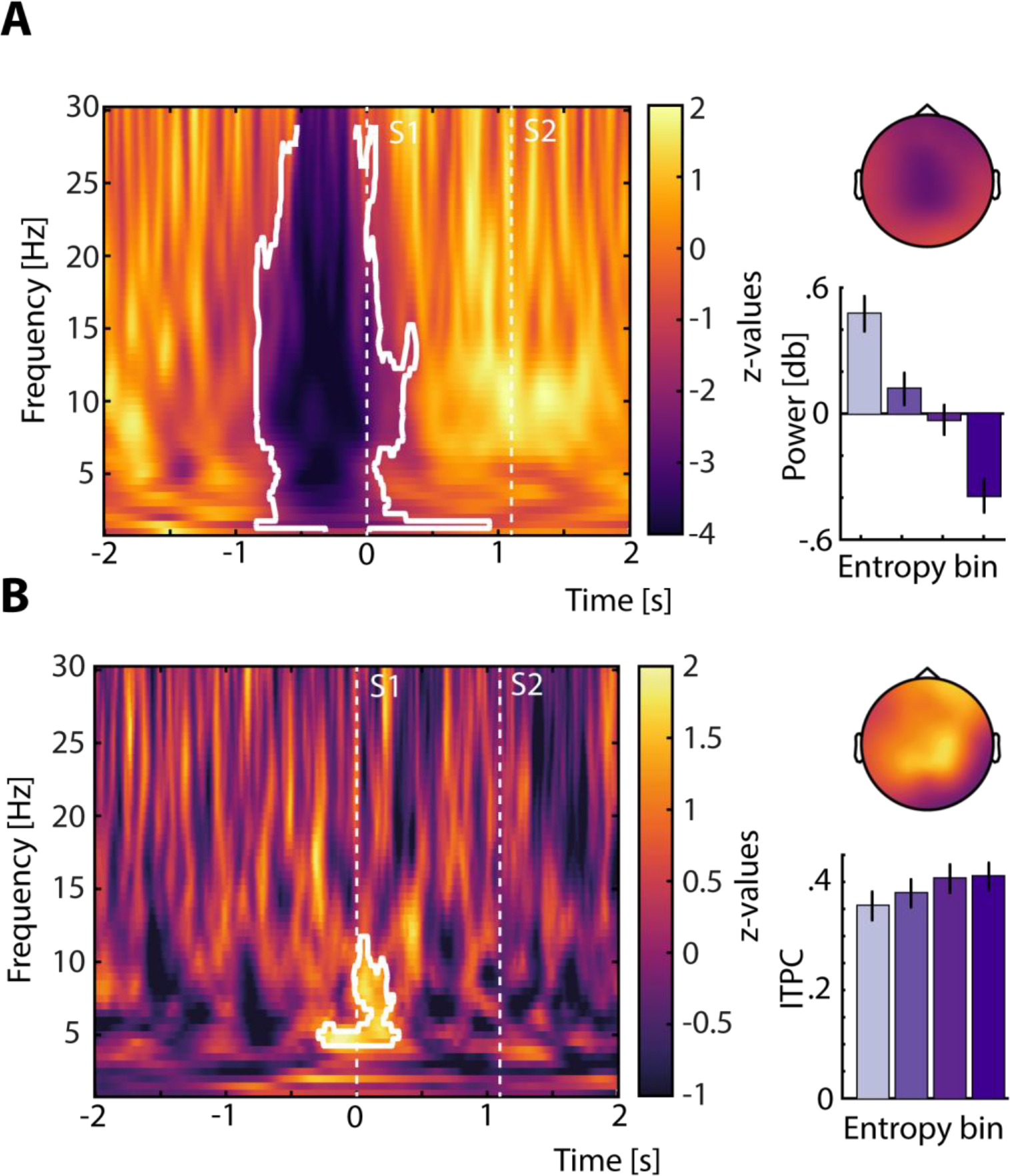
Time-frequency patterns of oscillatory power and phase coherence as a function of pre-stimulus entropy. A, Pre-stimulus power decreases with increasing entropy. Oscillatory power between 1 and 29 Hz significantly decreased with increasing pre-stimulus WPE. Colors indicate z-values resulting from a cluster-based permutation dependent samples t-test that compared slopes of linear trends across four bins of oscillatory power against zero, which were derived by sorting for increasing pre-stimulus entropy. The resulting negative cluster (*p* = .0009) is outlined in white and revealed a fronto-central topography. The bar graph shows average power within the cluster per bin (± 1 between-subject standard error of the mean, SEM). Darker colors correspond to higher pre-stimulus entropy. **B, Low-frequency phase-coherence at S1 increases with entropy.** ITPC around the onset of S1 increased with pre-stimulus entropy. Colors again indicate z-values from a similar cluster-based test as outlined for power, testing linear slopes across four bins of ITPC against zero. One positive cluster around S1 onset (–.25 to .30 s, 4–12Hz) with a predominantly central and frontal topography was found (*p* = .02) and is outlined in white. The bar graph shows average ITPC within the positive cluster per bin (± 1 SEM). Darker colors correspond to higher pre-stimulus entropy.

When analyzing inter-trial phase coherence as a function of binned pre-stimulus entropy, a later, positive cluster (–.25 to .30 s, 4–12 Hz) emerged around the onset of S1, indicating higher ITPC with increasing levels of pre-stimulus entropy (Fig. 3B, *p* = .02, *R* = .62) over a set of frontal and central electrodes. Accordingly, with increasing signal irregularity prior to S1, the neural response to S1 in a theta-/alpha-band became more phase-coherent.

## Pre-stimulus irregularity and peri-stimulus theta-phase angle predict perceptual decisions

Lastly, we explored the potency of our neural-irregularity measure, as well as the ensuing change in phase coherence during stimulus encoding, to explain the perceptual decisions based on the (physically identical) stimuli. Note in this context that neither ERP amplitude nor oscillatory power predicted decisions (see Fig. S6).

Sorting trials separately for each measure (see *Predicting perceptual decisions),* we then analyzed how the probability that a participant would choose the first tone as higher in pitch in a given trial would change, given this trial’s pre-stimulus entropy on the one hand, or this trial’s “phase similarity” at S1 onset (4–6 Hz,.01–.25 s post S1 onset, electrode Cz). Phase similarity is a single-trial measure we derived based on the observed (across-trials) ITPC effect. Phase similarity is calculated as the angular difference between a single trial phase at a given time and frequency (here:shortly after S1 in the theta frequency range) and the circular mean phase angle across trials. It thus captures how much a trial is deviating from the observed across-trials phase coherence in the time window of stimulus encoding.

Again, sorted trials were divided into four bins of equal trial number, where pre-stimulus entropy increased and angular distance from the mean angle decreased (i.e. ITPC increased) with bin number (Fig. 4). Linear slopes fitted to the legit-transformed probability of choosing S1 across bins were significantly positive for both pre-stimulus entropy (*t*_18_ = 3.15, *p* = .006, *r_e_*= .60) and angular distance from the mean (*t*_18_ = 3.20, *p* = .005, *r_e_* = .60). Note that sorting for pre-stimulus low frequency power did not result in significantly positive slopes (*t*_18_ = 1.7, *p* = .1, *r_e_* = .38; Fig. S6). Hence, a participant was more likely to perceive and decide on the first tone as higher in pitch if this first tone was presented in a period of relatively high neural irregularity. On the other hand, a participant showed a higher probability to decide for the first tone if the theta phase angle on that trial was more similar to the average theta phase angle across trials.

**Figure 4.**
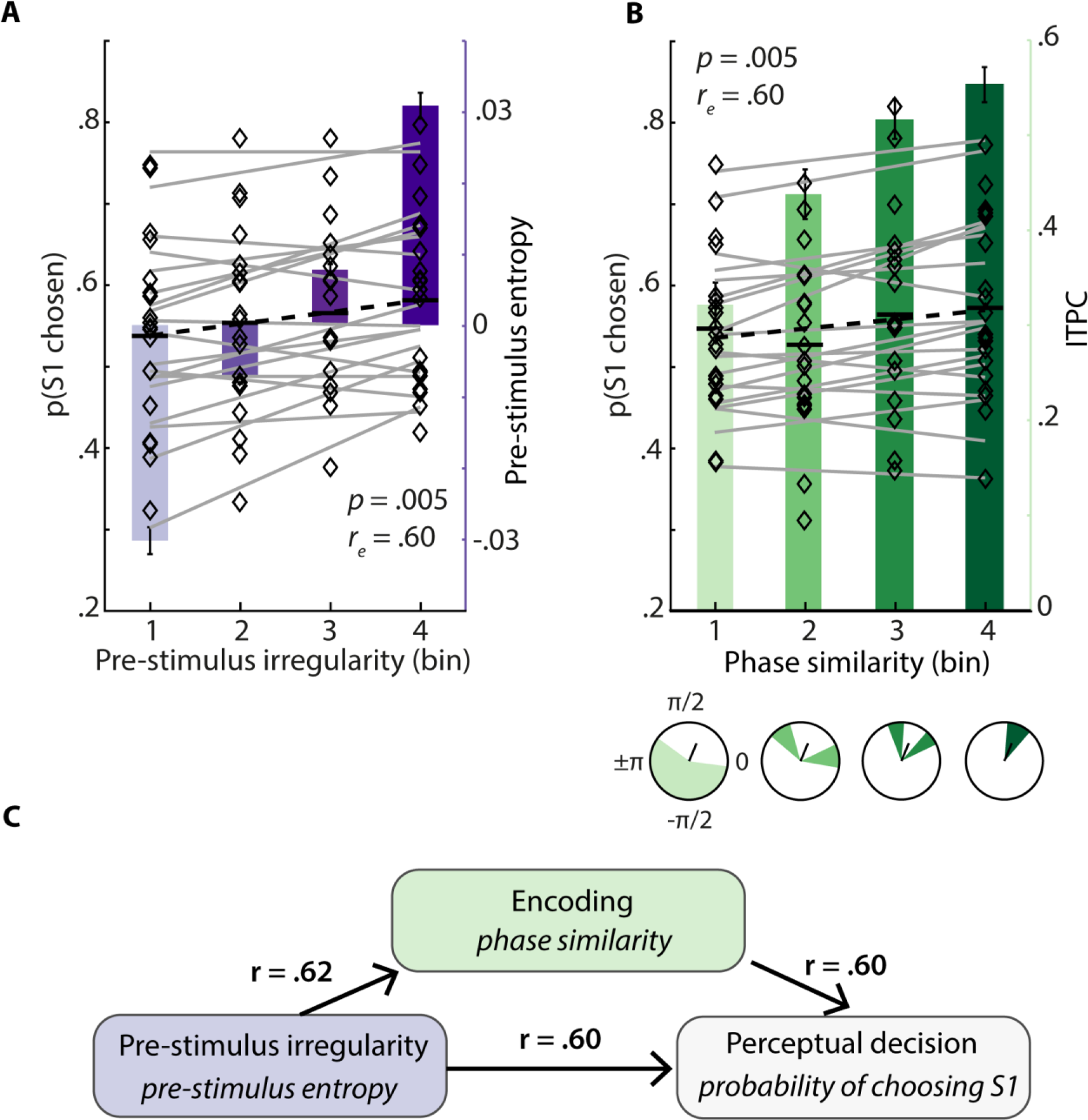
Pre-stimulus entropy and theta phase angle at sensory encoding both predict perceptual decisions. **A**, The probability of choosing S1 is shown for each subject and entropy bin (diamonds). Grey lines depict single-subject slopes fitted to probabilities across bins for visualization. Slopes were significantly positive on average (black dashed line, *p* = .005) representing an increase of choices for Sl with increasing pre-stimulus entropy (represented by bars of darkening lilac ± 1 SEM). **B**, Again probabilities of choosing S1 are shown per participant and bin together with grey lines representing slopes. Here, with increasing bin number, theta phase similarity at S1 onset increased, hence, ITPC increased (bars of darkening green ± 1 SEM). With increasing phase similarity, decisions for S1 got more likely (black dashed line, *p* = .005). Circle plots show distributions of phase angles per bins for an exemplary participant along with the mean angle across trials. **C, Links of irregularity, encoding and decisions.** The found relations of pre-stimulus irregularity with encoding and perceptual decisions as well as the link of encoding and perceptual decisions are visualized. Arrows imply timely, not directional connections. Note that *r* denotes the observed effect size. Colors correspond to measures as visualized in **A** and **B.**

## Discussion

Perceptual decisions in the absence of evidence should be driven by fluctuations in relevant states of the brain - that is, by the irregularity of ongoing neural activity. At the same time, such neural irregularity might be key to delineate neural changes that come with age. Here, we thus tested the potency of irregularity in the human EEG to describe both neural traits and states by (i) tracking age related changes in ongoing brain activity and (ii) by predicting discriminative decisions between two identical tones. Results can be summarized as follows: First, average irregularity of the EEG signal increased with age, paralleled by a reduction in variability. Second, individual pre-stimulus irregularity did not only capture low-frequency oscillatory power and influenced the encoding fidelity of presented tones but also biased perceptual decisions.

## EEG irregularity increases with age

Aging has been shown to manifest in the brain signal by lower temporal variability of hemodynamic activity (e.g. Garrett et al., 2013) and changes to the power spectrum of the EEG (e.g. Leenders et al., 2016), both possibly describe consequences of structural and functional dedifferentiation. But how does irregularity of the EEG signal relate to aging and the established markers of changing activity patterns?

Strikingly, the observed age-related increase of irregularity was accompanied by a decrease in the variability of EEG irregularity across trials (Fig. 1B). Note that this effect was not linked to previously observed age-related alterations in amplitude and latency of evoked responses (Pfefferbaum et al. 1980; Herrmann et al. 2016), since it was equally present both in pre-stimulus and whole-trial time windows (see Fig. 1B and C). Beyond offering a clear marker of aging in the EEG signal, these results close an important gap:they allow us to link electrophysiology to findings from fMRI research that emphasized reduced trial-to-trial brain signal variability with increasing age (Grady and Garrett 2014; Garrett et al. 2013).

Further validating our results, older participants showed shallower power spectral density (PSD) slopes. Both measures, entropy and PSD slope, are likely tracking the described twofold shift in the power spectrum that is also referred to as increased 1/f noise (Voytek et al. 2015). This, alongside the absence of ERP differences between age groups, renders it likely that the observed increase in entropy is in fact a time-domain reflection of a relative enhancement in higher-frequency noise in the signal. This interpretation gains further support from a comparison of weighted permutation entropy as used here with multiscale entropy approaches (Fig. S7; Mcintosh et al., 2014; Voytek and Knight, 2015).

If so, which structural and functional changes mediate these age-related alterations in brain signal variability and 1/f noise? During aging, the loss in number of neurons is negligible (Morrison and Hof, 1997), especially compared to the age-accompanying damage of white matter integrity (Sullivan, Rohlfing, and Pfefferbaum 2010), which among other modifications is responsible for a decline of large-scale structural networks. Importantly, the connectivity of higher-order as well as sensory functional networks shrinks with age (Andrews-Hanna et al. 2007; Geerligs et al. 2015). A shallower power spectrum thus might not only reflect higher degrees of 1/f noise (Voytek et al. 2015; Leenders et al. 2016) but also a decreased utilization of long-range neural networks (Meunier et al. 2009; Wang et al. 2010) and a parallel increase in local processing (Sleimen-Malkoun et al. 2015). Increased irregularity in older brains during an auditory discrimination task therefore, by proxy, also points towards reduced connectivity in functional networks. Furthermore, decreased trial-to-trial variability might trace back to reduced spatial variations of brain activity that has been shown to come with age (Garrett et al. 2013).

Additionally, age-related changes in the slope of power spectra over higher frequencies (30–70 Hz; Fig. S4) have recently been hypothesized to dynamically track changes in the excitation­ inhibition (E/1) balance via a positive relationship between PSD slopes and E/1 ratio (Gao, Peterson, and Voytek 2016). Older brains thus might show increased E/1 ratios as compared to younger ones. If that is the case, irregularity of the EEG possibly captures global inter-individual changes in the brain signal, that is, it describes age-related alteration as a neural trait. By design and as outlined in more detail below, but unlike PSD, entropy additionally offers information regarding ongoing activity on a trial-wise, time-resolved basis and thus allows estimating neural states.

## EEG irregularity tracks neural states and affects encoding

Non-human animal physiology research on the neural dynamics of sensory processing has recognized low-frequency power as a proxy of synchronized and desynchronized neural states in ongoing cortical activity (Marguet and Harris 2011; Pachitariu et al. 2015; Curto et al. 2009). Periods of relatively low power are used to indicate desynchronized states. A sensitive (and potentially more versatile) measure of neural irregularity or desynchronization should thus transfer these findings to the human brain and exhibit a negative relationship with low-frequency power. Notably, spontaneous variations in oscillatory power have also been linked to perceptual processes in humans (e.g. Boncompte et al., 2016; Kayser et al., 2016). Here, with higher irregularity prior to stimulus onset, power in low frequencies decreased. Although the frequency range may seem rather broad, decreases were strongest between 1 and 20Hz, a frequency range previously used as a proxy for neural states (Pachitariu et al. 2015).

The fronto-central topography of oscillatory power decreases appears compatible with auditory cortical generators and thus represents growing evidence for an impact of pre-stimulus neural state on the processing of sensory information. Unifying results from different methodologies and species, the entropy measure used in the present study (WPE) thus offers a new and promising approximation of irregularity in neural activity and thereby quantifies the degree of desynchronization in the human EEG.

Importantly, increasing pre-stimulus irregularity additionally was associated with higher phase coherence during the period of stimulus encoding. This observation aligns well with recent results from non-human animal work that related stronger desynchronization of ongoing activity to lower variability of neural responses to sensory stimulation (Schölvinck et al. 2015).

Furthermore, although a dependence of sensory encoding and cortical representations on the degree of neural irregularity has been noted before (Marguet and Harris 2011; Pachitariu et al. 2015; Curto et al. 2009), this relationship has not yet been revealed for humans. Neurally encoding the same stimulus in a more desynchronized state of cortical activity should lead to a more thorough sensory representation (e.g. Marguet and Harris, 2011). The present data propose that pre-stimulus irregularity influences the encoding and subsequent representation of the first tone (Snresulting in a higher-fidelity representation of S1 compared to S2.

This, in turn, should manifest in behavior: When comparing two physically identical tones, the tone that is represented more thoroughly should be more likely to be chosen, due to a higher fidelity or saliency with which it was neurally encoded (Fig. 4). The next section will discuss this in more detail.

## Pre-stimulus irregularity and encoding bias decisions

Decisions in the face of perceptual ambiguity are known to be influenced by pre-stimulus patterns in low-frequency power and phase of oscillations in the human EEG (e.g., Busch et al., 2009; Strauß et al., 2015; Wilsch et al., 2015). The impact of irregularity in ongoing activity on perceptual processes and decisions, however remains unknown. In addition to irregularity-driven changes in encoding, we find that with neural irregularity prior to S1 onset, S1 became more likely to be chosen as higher in pitch (Fig. 4A). Note that in this analysis we focused solely on intra-individual variations, accordingly comparing states as opposed to traits. Since the degree of irregularity in the EEG signal approximated the neural state of ongoing activity in terms of pre-stimulus desynchronization, we take this as evidence of the neural state influencing perceptual decisions. Thus, pre-stimulus irregularity state influences pending decision, likely by changing response bias (Limbach and Corballis 2016).

In addition, however, pre-stimulus neural irregularity also shaped neural encoding of to be judged stimuli, as evidenced by our ITPC analysis (Fig. 3). We would thus expect that this increased sensory­ encoding strength should also be directly predictive of perceptual decisions. Indeed, with increasing theta phase similarity at S1 onset, which can be thought of as a measure of optimized encoding, the probability that S1 was chosen also increased (Fig. 4B). This suggests an additional influence of the fidelity of early sensory representations on perceptual decisions (compare Fig. 4A and B). Again, a more thorough encoding of the first tone should lead to a more detailed and hence more salient representation, which gets chosen in the end of the trial.

The present data propose that, states of pre-stimulus irregularity influence perceptual decisions via changes to neural encoding which in turn exerts a separate impact on the decision via modulating the salience of representations (cf. Fig. 4C). We hereby expand research on the influence of spontaneous variations in brain activity on perceptual decisions in two different ways: First, we establish a trial-wise measure of irregularity in ongoing brain activity as compared to most studies of the field that relied on averaged activity estimates (e.g. Hesselmann et al., 2008; Amitay et al., 2013). Second, we here show trial-wise irregularity of spontaneous brain activity to indeed predict both the encoding of presented tones and the ensuing perceptual decision that follows at the end of each trial, as had been previously implied (Micheyl, McDermott, and Oxenham 2009; Bernasconi et al. 2011).

In sum, neural irregularity not only characterizes behaviorally relevant brain states, but also captures trait-like changes that come with age.

## Materials and methods

### Participants

Nineteen healthy participants (19–74 years; Median age 49 ± 18.75 siQR, 12 female) with self-reported normal hearing took part in the experiment. Participants gave written informed consent and were financially compensated. None of the participants reported a history of neurological or otological disease. The study was approved by the local ethics committee of the University of Lübeck.

### Stimulus material

One 650-Hz, 150-ms pure tone (sampled at 44.1 KHz, rise and fall times of 10 ms) was generated using custom MATLAB^®^ code (R2014b; MathWorks, Inc., Natick, MA). In each trial during the main experiment, this pure tone was presented twice (i.e., as a pair) with an inter­ stimulus-interval (lSI) of 900 ms using Sennheiser^®^ HD-25–1 headphones. To ensure accurate timing, auditory stimuli were presented via the Psychophysics toolbox (Brainard 1997) and an external low­ latency audio interface (Native Instruments, Berlin, Germany). Note that the tone pairs were presented perfectly audible (i.e., no masking noise) at a comfortable loudness level, which participants could adjust themselves during practice trials.

### General procedure

Participants were seated in a quiet room in front of a computer screen. First, after 20 practice trials, they underwent an adaptive tracking procedure that resembled the main experiment but consisted of pairs of different tones which gradually were rendered more similar (see Supplementary methods). Subsequently, they performed the main task, which consisted of identical stimuli only before carrying out a second run of the adaptive tracking procedure. In all parts of this study (practice trials, adaptive tracking, main experiment), participants had the task to decide on each trial which one of the two tones was higher in pitch.

### 2AFC pitch discrimination task

Subjects were instructed to indicate after each pair of presented pure tones, which one was higher in pitch by pressing one button on the computer keyboard. They were encouraged to include the degree to which the respective tone was perceived as higher in pitch by using a rating between 1 and 6 (Fig. 1). Here, a rating of 1 corresponded to the perception of “the first tone being clearly higher in pitch” and a rating of 6 to “the second tone being clearly higher in pitch”. The mapping of response buttons was reversed for approximately half of the subject sample (9 of 19 subjects, in random order).

A first 150-ms tone (S1) was followed by a fixed 900-ms silent interval before the second 150-ms tone (S2) was presented. Subsequently, the response screen was shown until subjects entered their response with a limit of 2 s (Fig. 1). A grey fixation cross was presented throughout the trial and subjects were instructed to fixate their vision on it at all times. All visual content was presented and responses were recorded using custom MATLAB^®^ scripts and the Cogent 2000 toolbox.

After an inter-trial-interval, randomly jittered between 2 and 4 s, the next trial started, indicated by the fixation cross changing its color from grey to light green and back to grey within 500 ms. The experiment consisted of a total of 500 trials, divided over five blocks of 100 trials each. The first trial of each block was started by the participants as soon as the experimenter had left the room. Blocks were separated by short breaks. The experiment took about 60 minutes in total.

Bogus feedback was provided after each of the first 20 trials in every block, where, in 65% of all feedback trials, a positive feedback indicating a correct pitch discrimination was given. This proportion of positive bogus-feedback was chosen to keep participants engaged in the task and took into account a comparable previous research design (Bernasconi et al. 2011). In the case of trials involving auditory bogus feedback, the response screen was followed by a sound indicating correct or incorrect answers after 100 ms. Additionally, after every 20th trial, sham accuracy scores representing average performance in the past 20 trials, randomly chosen from a uniform [S5; 65]-% distribution were displayed on the screen for 3 s. Note that, for further analyses, all trials followed by feedback were excluded.

After the experiment, subjects frequently reported that they perceived pronounced pitch differences between the two (physically identical) tones. Before debriefing, no subject raised concerns regarding the actual nature of the stimuli. In fact, behavioral pilot experiments (n = 10) showed that even participants who knew that a high proportion of all trials would contain identical stimuli still reported to perceive considerable differences in pitch. In combination with previous experimental work that used identical stimuli in a comparable task structure (Amitay, Irwin, and Moore 2006; Amitay et al. 2013; Bernasconi et al. 2011) we are thus confident that participants paid attention to the task and engaged in it. Additionally, a computational modelling study that estimated the magnitude of perceived differences in similar tasks as pronounced as 8.5 Hz in 50% of all trials and larger than 20 Hz in 10 % of all trials (Micheyl, McDermott, and Oxenham 2009) supports the feasibility of such a task structure.

### Behavioral data analysis

As expected with physically identical stimuli, only few participants exhaustively used the full range of “first vs. second tone had higher pitch” ratings (1–6). Most ratings were instead centered on the middle range (3–4) of possible ratings. The variance of responses over trials showed no correlation with participant age (*r* = –.07, *p* = .78). We thus binarized the rating responses and coded ratings 1–3 as “first stimulus chosen” and ratings 4–6 as “second stimulus chosen”. Proportions of both binarized response types were logit transformed (see *EEG time- and frequency domain analyses)* prior to statistical analysis to approximate normality for the originally [0; 1]-bound proportions. To check for response bias (i.e. favoring one choice over the other), these transformed proportions were compared using a paired t-test. We additionally tested for and excluded the presence of more complex response strategies by analyzing autocorrelated responses (see supplementary methods and Fig. S1).

### EEG recording and preprocessing

EEG was recorded with a 24-channel mobile EEG setup (SMARTING, mBrainTrain, Belgrade, Serbia) at a sampling rate of 500Hz, a resolution of 24 bits and a bandwidth of 0–250 Hz. Electrode impedances were kept below 10 kΩ. The amplifier was attached to the EEG cap (Easycap, Herrsching, Germany). Recordings were online referenced to electrode FCz and grounded to electrode AFz. All data and event-related triggers were transmitted to a nearby computer via Bluetooth where both were saved using the Labrecorder software, which is part of the Lab Streaming Layer (LSL; Kothe, 2014).

Offline, continuous data were bandpass filtered from 0.5 Hz (−12 dB attenuation at 0.03 Hz) to 100Hz (−12 dB attenuation at 100.2 Hz) with a zero-phase finite impulse response filter (filter order 1200). Filtering and all other analysis steps were carried out in MATLAB^®^ 2014b using custom scripts and the fieldtrip toolbox (Oostenveld et al. 2011). After re-referencing to average mastoids, we epoched the data from −2 s to +2 s around onset of S1. The first 20 epochs of every block were excluded from further analysis since they involved feedback. An independent component analysis was carried out and components related to eye-movements, muscle activity or heartbeat were identified and removed from the data on the basis of their time- courses, frequency spectra and topographies. Subsequently, remaining artefactual epochs were removed after visual inspection. On average, 43 % (± 12% SD) of components and 7% (± 4% SD) of trials were removed.

### Information theoretic measure

Entropy is taken to quantify the information contained in a signal. Accordingly, a variety of entropy-based measures such as Shannon Entropy (Shannon 1948), Kolmogorov-Sinai Entropy (Kolmogorov 1968), Approximate Entropy (Pincus 1991) or Sample Entropy (Richman and Moorman 2000) has been suggested and has been used to estimate the complexity of natural time series data. Each of them however is constrained by required metric properties of the input data. In contrast, Permutation Entropy (PE) is a form of symbolic or rank-based entropy measure (Bandt and Pompe 2002) that estimates the complexity of a time series while being robust against various degrees of non-linearity in the signal. Furthermore, PE takes into account the temporal structure of a signal and is computationally very efficient (Riedl, Müller, and Wessel 2013). Permutation entropy is thus particularly attractive as an entropy measure for biological time-series as obtained with EEG, since these incorporate non-linear processes at least to some degree. Although a full introduction on the principles of entropy calculation is beyond the scope of this article, the basic rationale of PE (and, as used here, a weighted version of PE; WPE) shall be outlined briefly below.

The basic calculation of PE can be split up in three steps: First, data are divided into overlapping windows of reasonable size (e.g. 50 samples, but see Staniek and Lehnertz, 2007 for an overview of parameter selection for PE). Second, the time series data within the windows are mapped into symbolic space by dividing each window into short sub-sequences (e.g. 3–5 samples size, 1 sample distance) and calculating their ranks, resulting in so called motifs. Importantly, the number of different motifs that can occur is determined by the number of samples in one sub­ sequence (motif length). Using a motif length of 3 for example results in 3! = 6 possible motifs. Third, the occurrences of each motif within each bigger time window are counted and their frequency of occurrence is used to calculate one Shannon Entropy value for each motif. These values finally are added before being multiplied with −1 to arrive at the PE of one window. Given a sufficiently high sampling frequency (i.e., if the number of samples within a window allows all possible motifs to occur more than once), PE can be calculated with high precision, still offering satisfying time resolution.

The defining feature of PE as compared to other entropy algorithms is that the calculation of PE is not based on the real values of the time series but on sequences of ranks, the so-called motifs. While this makes PE very robust against outliers and suitable for non-linear time-series, it disregards, by design, all metric information in a given data window. For example two windows containing the voltage (l.lV) measurements [−1, +1, +4] and [−20, +60, +150] would be mapped onto the same motif (1, 2, 3). To overcome this potential limitation, Fadlallah et al. (2013) proposed a weighting of occurrence frequencies that enter PE calculation:The relative frequency of occurrence of every motif is weighted by the variance of the data segments from which this motif was derived, so that its contribution to the resulting Weighted Permutation Entropy (WPE) is increased for increasing variance. As a result, WPE retains all features of PE but is more sensitive to abrupt changes in a signal as revealed by the application to both synthetic noisy time series and EEG data (Fadlallah et al.2013).

We calculated WPE (in the following and above referred to as entropy) following the same procedure as outlined above for the extraction of PE time courses: For each electrode and trial, entropy was computed using a sliding window of 50 samples (corresponding to 100 ms in the present data) that was shifted in steps of 10 sampies (20 ms), a motif length of 3 and a time delay fa ctor of 1. This resulted in entropy time-courses of 200 samples per trial.

To explore the impact of pre-stimulus entropy on low-frequency power fluctuations, encoding and perception of the presented tones, trials were sorted for increasing baseline-corrected pre-stimulus entropy (averaged over the interval from –.4 to –.1 s at electrode Cz). Subsequently, sorted trials were separated into four equally sized, non-overlapping bins (number of trials / 4) resulting in two grouping variables per trial: the given response and the bin membership, whereas for the latter higher values corresponded to higher levels of pre-stimulus neural irregularity.

### EEG-irregularity as an inter-individual marker of aging

To compare the degree of irregularity between participants of different age, entropy at every electrode was averaged per subject over all trials and subsequently correlated with age. Since trial-wise averages of entropy did not only display considerable inter- but also intra-individual variance, which appeared to shrink with increasing age (see Fig. 1B), we calculated the intraclass correlation coefficient (ICC). Note that the relationship of entropy and age was strikingly unaffected by different time-windows that were used for averaging before correlating averages with age (Fig. 1B, but see Fig. S3). Since our primary interest lay in the irregularity patterns of ongoing activity, we used pre-stimulus entropy for all further analysis.

To relate the fairly recent measure of entropy to other, commonly suggested measures of neural irregularity (Leenders et al. 2016; Voytek et al. 2015), we also calculated the power spectral density (PSD) from subjects’ trial data using 2-s-windows (which overlapped by 50%, yielding three PSD estimates per trial that subsequently were averaged). An individual linear fit across frequencies between 1 and 30Hz (excluding the 8–13-Hz alpha range; Voytek et al., 2015) was obtained for the resulting PSD estimate. The slope of the resulting line can be used to estimate the degree of 1/f noise in the power spectrum (Bédard, Kröger, and Destexhe 2006; Berthouze, James, and Farmer 2010). In brief, a more uniform distribution of neural oscillatory power across frequencies (i.e., a shallower spectrum) is indicative of increasing neural irregularity. We thus expected flatter spectra and accordingly more positive regression slopes for older compared with younger participants. To test this, we calculated the Pearson correlation of age and the slope of the linear regression Iine fitted to the power spectrum.

### Basic EEG time- and time-frequency domain analyses

EEG data were separated into trials for which a response indicating the first versus second tone was perceived as higher in pitch. Time-locked averages (i.e., event-related potentials, ERPs) were computed for every participant and condition (first tone chosen vs. second tone chosen) separately.

To assess potential differences in ERP amplitude on the group level, we performed a cluster­ based permutation dependent samples t-test (Maris and Oostenveld 2007), comparing the ERP in trials where the first versus the second tone was later chosen as higher in pitch. This test proceeds by first clustering adjacent bins with t-values with p < .05 in electrode-time space, where a cluster consisted of at least three neighboring electrodes. The sum of all t-values in a cluster subsequently was compared against the distribution of 10, 000 random clusters, which were generated by iteratively permuting the labels of conditions. The resulting p-value of a cluster corresponds to the proportion of performed Monte Carlo iterations that exceed the summed t-values of the empirically observed cluster (Maris and Oostenveld, 2007; for an application see e.g., Wöstmann et al., 2015).

Complex-valued time-frequency Fourier representations of the data were obtained by means of convoluting single-trial time courses with frequency-adaptive Hann-tapers of 4 cycles width. Oscillatory power from 1 to 30 Hz (in .5 Hz steps) and from 30 to 100 Hz (2 Hz steps) was obtained by squaring the modulus of the Fourier spectrum. The average power estimate across trials was then baseline corrected and represented as change in Decibel (dB) relative to the mean power during a [−2; −1] s pre-stimulus time-window. Note that this procedure was performed separately for each of the four bins that resulted from sorting for pre-stimulus entropy as outlined above. To relate average oscillatory power to the pre-stimulus state of the brain (i.e. the level of irregularity), we performed a subject-wise linear fit across all bins, separately for every electrode, time-point and frequency. The resulting slope qualifies the relationship of pre-stimulus irregularity and oscillatory power. Slopes were tested against zero using the same cluster-based approach outlined above for time-domain data.

### Inter-trial phase coherence (ITPC)

We then calculated Inter-trial phase coherence (ITPC; 0 ≤ ITPC; ≥ 1) for every bin of sorted trials to test for differences in phase coherence between different levels of pre-stimulus neural irregularity. Note that ITPC can also be expressed as 1 - circular variance (e.g. Breakspear et al., 2010), where high ITPC values reflect relatively low levels of variance in the precise phase angle across trials at a given time (e.g. stimulus onset) and frequency. Thus, ITPC poses an across-trials proxy for the variability of phase angles. Since measures of phase coherence are strongly influenced by the number of trials on which they are computed (Ding and Simon 2013) we used the same number of trials for each participant from four levels of pre-stimulus irregularity (see *Information-theoretic measure*) to calculate ITPC and power estimates. For further metric and statistical analysis, the [0;1]-bound ITPC measure at every electrode e, time-point t and frequency f was logit-transformed, an approach previously shown to outperform other transformation techniques for binomial and proportional data (Warton and Hui 2011). This resulted in a [−∞; +∞]- bound measure of

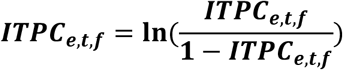

To analyze how pre-stimulus irregularity influences phase-coherent responses to the first tone, for each subject, linear fits across the four ITPC estimates were performed per electrode, timepoint and frequency as described above for average oscillatory power. Here, slopes approximate the relation of pre-stimulus entropy and the consistency of phase angles in response to a stimulus and were tested against zero using the same cluster-based approach outlined above.

### Predicting perceptual decisions

As outlined above, trials were sorted ascendingly for pre-stimulus entropy as a marker of irregularity in the EEG-signal before stimulation onset. Importantly, we did not only expect pre-stimulus irregularity to impact the encoding of presented stimuli but ultimately to affect pitch perception and thereby the impending decision. The sorting of trials for pre-stimulus entropy thus also was used to establish bins of behavioral responses.

Additionally to the influence of pre-stimulus state on encoding that was revealed by a positive cluster around stimulus onset (Fig. 2), the encoding of a tone and the ensuing decision might have been influenced by variations in evoked responses unrelated to spontaneous changes in the pre-stimulus state. Thus, for every trial, the phase angle at the central point of the found cluster (4–6 Hz, .01 – .25 s, electrode Cz) was extracted and compared to the corresponding average phase angle across trials by computing the absolute angular distance, resulting in “phase similarity”. The result, one value of phase similarity per trial, captures the trial-wise difference in phase from the stereotypic response during encoding. Subsequently and analogous to the approach used for pre­ stimulus entropy, trials were sorted for increasing phase similarity and split up into four non­ overlapping bins of equal trial number. Hence, to the existing grouping variables (response, entropy bin) per trial, another one, phase similarity bin membership, was added. Note that phase-angles were more similar to the average phase with increasing bin-number and hence ITPC can be expected to increase with bin-number.

To quantify the impact of pre-stimulus irregularity and evoked responses on the ensuing perceptual decision, the probability of choosing the first tone was calculated for each participant and bin before undergoing logit transformation. Per subject and binning variable (entropy / phase similarity), a first order polynomial was fitted to the legit-transformed probabilities across all four bins. The resulting slopes were tested against zero using a one-sample t-test.

### Effect sizes

As a measure of effect size for t-statistics resulting from both dependent- and independent-sample t-tests, we calculated the *r_equivalent_* (Rosenthal and Rubin 2003) henceforth denoted as *r_e_*. For multiple comparisons, e.g. when comparing electrode × frequency × time pairs, we averaged effect sizes of all t-tests within a significant cluster to estimate *R_e_* (Strauß et al. 2015; Wöstmann et al. 2015).

## Acknowledgements

Research was supported by the Volkswagen foundation (to JO; BIT-CHAT) and the European Research Council (to JO;ERC-2014-CoG AUDADAPT).

